# Proteomic mapping of macrophages in response to the clearance of apoptotic cells reveals a unique reprogramming profile

**DOI:** 10.1101/2023.11.23.567533

**Authors:** Benjamin Bernard Armando Raymond, Andrew Michael Frey, Matthias Trost

**Affiliations:** Laboratory for Biomedical Mass Spectrometry, Biosciences Institute, Newcastle University, Newcastle upon Tyne, UK

**Keywords:** Efferocytosis, Macrophage, SILAC, Inflammation, Data-independent acquisition

## Abstract

During the daily process of healthy cellular turnover, billions of cells undergo apoptosis in the human body. These cells are removed by phagocytic cells, namely macrophages through a process known as efferocytosis, which triggers a cascade of reprogramming events in the cell, with a shift towards a pro-resolving or ‘wound healing’ phenotype. To date, no study has attempted to investigate these phenotypic changes from a proteomic perspective. Here, we present a novel and robust workflow for the investigation of proteome and secretome changes in bone marrow-derived macrophages (BMDMs) following efferocytosis using stable isotope labelling by amino acids in cell culture (SILAC) combined with data-independent acquisition (DIA) mass spectrometry. We show that using this workflow we can dissect the mixed proteomes of BMDMs and apoptotic cells to specifically map the reprogramming events occurring in macrophages in the later stages of efferocytosis. Specifically, we find that efferocytic macrophages adopt an alternatively activated phenotype underpinned by an increase in efferocytic and anti-inflammatory markers. We also show that the secretome contains factors that can reprogram naïve BMDMs towards an efferocytosis-like, pro-resolving, phenotype. Our results provide an unprecedented view of the efferocytic landscape of macrophages and will aid in further understanding this important immunological process in the larger context of immune homeostasis and inflammatory disorders.

## Introduction

The clearance of dying cells by professional and non-professional phagocytic cells plays a fundamental role in tissue homeostasis and resolution of inflammation. This process, known as efferocytosis, is critical for maintaining immune tolerance and prevents the release of autoantigens from apoptotic cells that could trigger autoimmune responses. Consequently, dysregulation of efferocytosis has been implicated in various diseases, including autoimmune and inflammatory respiratory disorders^1^. Macrophages, one of the principal efferocytic cell types, express various pattern recognition receptors that are specialised in recognising antigens on dying cells, triggering phagocytosis. As key immune regulators, macrophages can modulate the immune response via their ability to shift between pro (M1)- and anti (M2)-inflammatory states and upon efferocytosis, they acquire an M2-like or pro-resolving phenotype in order to dampen the immune response and promote resolution of inflammation. Despite the importance of macrophage efferocytosis in health and disease, our understanding of the molecular mechanisms underlying this reprogramming process is limited. While studies have investigated transcriptomic changes in macrophages following the early stages of efferocytosis^2^, no study has investigated changes in the proteome in order to specifically map the phenotype of efferocytic cells. Notably, it has been shown that metabolites secreted by apoptotic cells^3^ and extracellular vesicles produced by efferocytic macrophages ^4^ can induce a pro-resolving phenotype in neighbouring cells and promote efferocytosis.

In this study, we sought to address this knowledge gap by employing a proteomic approach to investigate changes in bone marrow-derived macrophages (BMDMs) following phagocytosis of apoptotic Jurkat cells. Our aim was to gain a deeper understanding of the molecular mechanisms underlying the events following macrophage efferocytosis, which could elucidate novel targets for therapeutic intervention. By using stable isotope labelling by amino acids in cell culture (SILAC), we were able to separate the proteomes of BMDMs and Jurkat cells, and by combining this with data-independent acquisition (DIA) mass spectrometry, we achieved a deep level of proteome coverage. Our data show that, as expected, macrophages undergo extensive reprogramming following efferocytosis, shifting towards a unique phenotype that is, for the first time, mapped on the proteome level. Our data also profiles the secretome of efferocytic BMDMs, containing a mixture of macrophage- and apoptotic cell-derived proteins, that when added to naïve BMDMs, alters their phenotype to one resembling efferocytic BMDMs. This robust methodology has widespread applications for profiling efferocytic macrophages from different biological or disease contexts, such as those from inflammatory disorders, and thus, can be used to better understand how defective efferocytosis may contribute to disease.

## Results

### BMDMs undergo extensive proteome changes following efferocytosis

While the advent of label-free quantitative mass spectrometry has revolutionised the proteomics field, some limitations exist, such as being able to distinguish between mixed proteomes from closely related species. This has made the examination of proteome changes in macrophages following the phagocytosis of same- or similar species particularly challenging. To achieve this, we have implemented a novel approach by employing stable isotope labelling by amino acids in cell culture (SILAC) and combining this with data-independent mass spectrometry to improve quantification. We initially sought to establish an easy-to-reproduce model of efferocytosis using murine bone marrow-derived (BMDMs) and apoptotic Jurkat cells (Fig. 1A). Firstly, we were able to induce early apoptosis in ∼80% of Jurkat cells after 3 h treatment with the kinase inhibitor staurosporine, based on the percentage of Annexin V/7-AAD double positive events (Fig. S2). Using confocal microscopy, we could show that after ‘pulsing’ BMDMs with apoptotic Jurkat cells and ‘chasing’ for 30 min we could observe that apoptotic cells were fully engulfed within BMDMs and not merely stuck to them (Fig. 1B). We then employed flow cytometry to quantify the level of efferocytosis in BMDMs, demonstrating that ∼60% of BMDMs contained apoptotic cells, after a 30 min chase, which reduced to ∼40% after 3 h, likely due to dye quenching/degradation in the lysosomal compartment following phagocytosis (Fig. S3).

**Figure 1:**
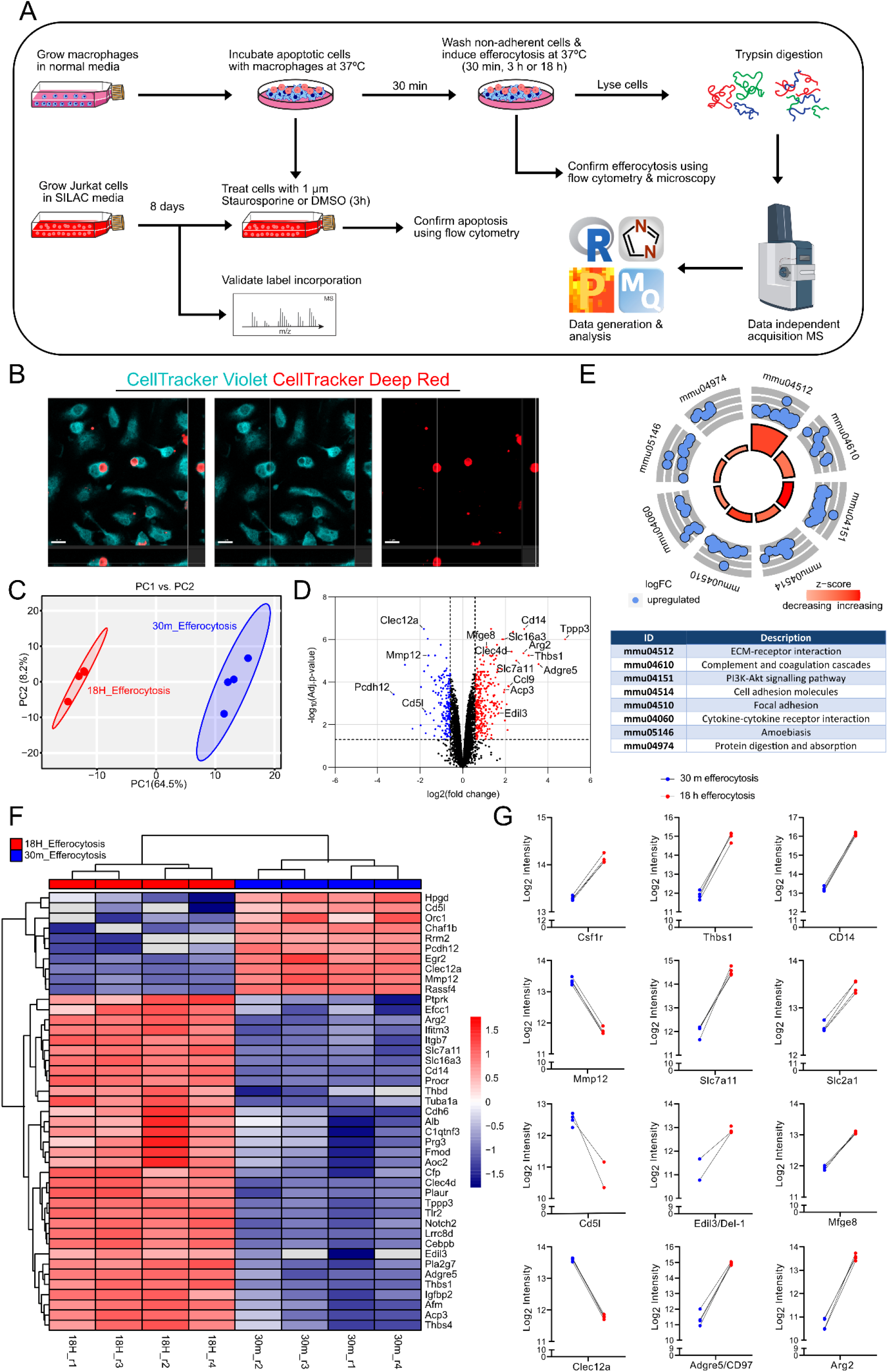
Macrophages undergo extensive proteome changes following efferocytosis. (A) Schematic of the workflow to examine proteomic changes in total cells following efferocytosis, from cell culture to data analysis. (B) Confocal microscopy orthogonal sections of CellTracker Violet-stained BMDMs phagocytosing CellTracker Deep Red-stained apoptotic Jurkat cells after 3 h. Orthogonal views show Jurkat cells being fully engulfed by macrophages. Scale bar is 10 μm. (C) Principal component analysis of 18 h and 30 min efferocytosis samples plotted as PC1 vs PC2. (D) Volcano plot showing differentially regulated proteins after 18 h of efferocytosis compared to 30 min. Fold-change cut-offs were set at > 1.5 and adjusted p-value cut-offs set at < 0.05. Data represents 4 biological replicates. (E) Gene ontology enrichment, using KEGG, of proteins with increased abundance following efferocytosis, shown as a circle plot. Increasing z-scores are shown in red, with the size of each bar representing significance based on the adjusted p-value, and log fold changes for each protein enriched within a specific term are expressed as blue bubbles. KEGG terms are described below in box. (F) Heatmap of the proteins with the greatest fold-changes 18 h following efferocytosis. Z-scores are shown on the right with increases shown in red and decreases shown in blue. (G) Selected proteins with fold changes greater than 1.5 and adjusted p-value < 0.05 that have known functions associated with efferocytosis or macrophage biology. Individual paired data points are shown.

For our proteomics-based efferocytosis assay, we incubated BMDMs with apoptotic Jurkat cells for chase times of 30 min or 18 h, where 30 min was treated as the control/baseline and 18 h as our efferocytosis condition, followed by lysis, trypsin digestion and mass spectrometry. Our initial observation of the non-normalised log2 transformed data showed that the mixture of light and heavy channels did not skew the light-channel across control and efferocytosis conditions (Fig. S4A). Post-filtering, we identified 6513 proteins that matched to *Mus musculus* in the light channel, and 4646 proteins that matched to *Homo sapiens* in the heavy channel (Fig. S4B). This indicates we could successfully differentiate between proteins from the light and heavy channels. Correlation matrices showed that inter-sample variance within each experimental condition was minimal (Fig. S4C), and a principal component analysis (PCA) demonstrated that replicates clustered together while experimental conditions had a good level of separation (Fig. 1C).

Following limma analysis of the murine proteins from the light channel, when comparing our 18 h efferocytosis condition to 30 min, we found 294 proteins with a fold-change > 1.5, and 152 proteins with a fold-change < −1.5 with an adjusted p-value < 0.05 (Fig. 1D). Gene ontology (GO) analysis by KEGG of proteins with increased abundances showed enrichment of pathways associated with extracellular matrix, complement, cell adhesion, cytokines, and the PI3K-Akt pathway, indicating a unique phenotype in BMDMs after undergoing efferocytosis, that has not been previously described (Fig. 1E). Further investigation into proteins that had the largest abundance changes following efferocytosis highlighted numerous proteins of interest (Fig. 1F). This included Adgre5, also known as CD97, which has been shown to antagonise pro-inflammatory activation of macrophages^5^ and Arginase 2, which like Arginase 1, promotes an anti-inflammatory phenotype in macrophages^6^ (Fig. 1F-G). Notably, we also identified numerous efferocytosis-related proteins, such as thrombospondin 1^7^, CD14^8^, Edil3^9^ and Mfge8^10^ (Fig. 1G). We also identified solute carrier transporters that have been previously described to play a role in macrophage polarisation as well as efferocytosis, such as Slc7a11^11^ and Slc2a1^2^. While Slc16a1/MCT1, a lactate transporter previously shown to be upregulated following efferocytosis^2^, was not found in our analysis, a closely related member of the monocarboxylate transporter family, Slc16a3/MCT4, was found to increase following efferocytosis (Fig. 1G). We also observed numerous proteins unique (and present in at least 3 of 4 replicates) to the efferocytosis conditions but not in any replicates of control conditions (and vice-versa) (Table S1). Despite not being able to calculate fold changes for these proteins, they may be playing a role in macrophage reprogramming following efferocytosis.

While gene ontology (GO) enrichment could not highlight any specific pathways or processes in proteins with decreased abundances, there were specific proteins of interest identified, such as Clec12a, a pro-inflammatory C-type lectin receptor whose expression on myeloid cells has been shown to correlate with disease activity in those with rheumatoid arthritis^12^, and Mmp12, a matrix metalloproteinase typically associated with pro-inflammatory macrophages^13^. Notably, the vast majority of decreased proteins (as well as those absent following efferocytosis, highlighted in Table S1) have roles associated with DNA-binding and cell cycle, suggesting that efferocytic BMDMs begin to resemble terminally differentiated, tissue resident macrophages, lacking the ability to proliferate.

Taken together, our data indicates that our SILAC workflow can successfully map reprogramming events in macrophages following efferocytosis. This has provided novel insights into this important immunological process, and highlights that macrophages shift to an alternative activated state upon apoptotic cell clearance.

### The efferocytic supernatant can alter the phenotype of naïve BMDMs

It has been shown that both macrophages and apoptotic cells secrete factors that promote a wound-healing phenotype in neighbouring cells to resolve inflammation^3,4^. In order to investigate this concept further using our SILAC workflow, we collected the supernatants from the BMDMs 6 h and 18 h following efferocytosis for mass spectrometry analysis. While PCA showed differences in how the same conditions clustered at the two timepoints (Fig. S5A-B), correlation plots demonstrated that samples were not largely different between the timepoints, and that intra-sample variability was low (Fig. S5C). In total, we identified 4355 light-labelled murine proteins across both timepoints and conditions, and after performing limma analysis found 145 and 1029 proteins with increased abundances at 6 and 18 h, respectively (Fig. S6A). Of those proteins that are shared between the timepoints ∼20% related to protein phosphorylation, based on GO enrichment of biological process (BP) terms (Fig. S6B-D). Notably, the same analysis of proteins with decreased abundance in the supernatants following efferocytosis at both timepoints highlighted proteins associated with cell adhesion and inflammatory responses (Fig. S6E-F). Thrombospondin-1 and Il1r2 had some of the most pronounced changes in their secreted abundance following 18 h efferocytosis, which is in agreeance with their putative roles as pro-resolving factors during apoptotic cell clearance^7,14^ and inflammatory conditions^15,16^.

Surprisingly, when analysing proteins from the heavy channel, i.e., from apoptotic Jurkat cells, we identified 3321 and 2794 proteins in the secretomes at 6 h and 18 h, respectively. However, due to the experimental design, fold-changes could not be calculated, and so protein intensities were log_10_ transformed and ranked. In the apoptotic cell secretome at both timepoints, largely the same proteins were ranked highest, including apolipoprotein E (ApoE), a known factor secreted by apoptotic cells that promotes their clearance^17^ and MIF, which is known to be packaged in apoptotic cell extracellular vesicles (EVs), and promotes macrophage proliferation during tissue maintenance^18^ (Fig. 2B). While it cannot be discounted that these proteins are being secreted from non-phagocytosed apoptotic cells, rigorous washing steps were performed during our pulse-chase assay which should have removed the majority of these cells. Therefore, we can hypothesise that these proteins arise from digested apoptotic cells that are exported from BMDMs via exocytosis. When further interrogating the human component of our secretome dataset, it was noted that GO analysis by cellular compartment showed a strong enrichment of EVs (Fig. S7A). While it was not within the scope of this study to characterise EVs following efferocytosis, it is interesting to speculate that this may be the case, as it has been recently shown that efferocytic macrophages secrete EVs^4^. Transmission electron microscopy (TEM) of cleared supernatants following ultracentrifugation did indeed show the presence of EVs in both control and efferocytosis conditions (Fig. S7C-D). Using our SILAC workflow, it will be pertinent to investigate the protein cargo of these EVs and their potential pro-resolving capabilities.

**Figure 2:**
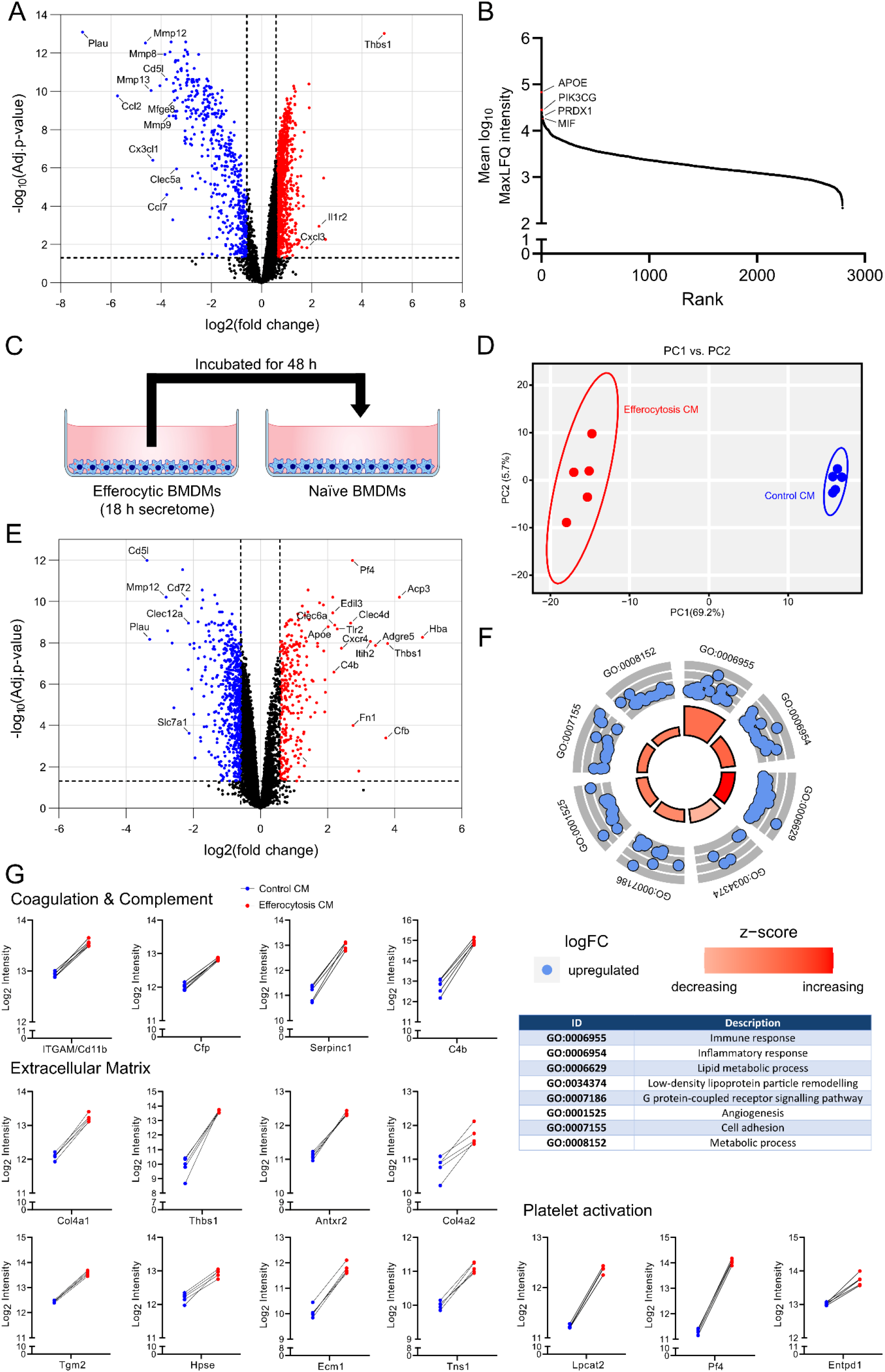
The efferocytic supernatant contains factors that can reprogram naïve BMDMs. (A) Volcano plot showing protein abundance changes of the BMDM secretome following 18 h efferocytosis. Data represents 5 biological replicates, except in the case of control experiments where one biological replicate was removed as it was an outlier. (B) Protein intensity ranking (log_10_) of heavy-labelled Jurkat proteins from 18 h efferocytosis supernatants. Intensities were calculated using MaxLFQ. (C) Schematic of workflow for creating conditioned mediums from control and efferocytosis experiments. Medium was collected, centrifuged to remove cells and debris, and incubated directly with naïve BMDMs for 48 h. (D) Principal component analysis of cCM- and eCM-treated BMDMs plotted as PC1 vs PC2. (E) Volcano plot showing protein abundance changes in BMDMs after treatment with eCM compared with cCM. Data represents 5 biological replicates. (F) Gene ontology (GO) enrichment, using Biological Process, of proteins with increased abundance 48 h following eCM treatment, shown as a circle plot. Increasing z-scores are shown in red, with the size of each bar representing significance based on the adjusted p-value, and log fold changes for each protein enriched within a specific term are expressed as blue bubbles. GO terms are described below in box. (G) Selected proteins with fold changes greater than 1.5 and adjusted p-value < 0.05 that have known functions associated with coagulation and complement, extracellular matrix, and platelet activation. Individual paired data points are shown. For volcano plots, fold-change cut-offs were set at > 1.5 and adjusted p-value cut-offs set at < 0.05.

We then sought to investigate whether or not these supernatants could alter the phenotype of naïve BMDMs. To do so, we incubated BMDMs with cleared, 18 h control (c) or efferocytic (e) supernatants, hereon referred to as conditioned medium (CM) for 48 h (Fig. 2C). PCA demonstrated that cells treated with eCM clustered separately from those treated with cCM (Fig. 2D), and we identified 275 and 556 proteins with increased and decreased abundances after eCM treatment, respectively (Fig. 2E). Gene ontology enrichment analysis of proteins with increased abundance identified pathways associated with the immune response, lipid metabolic process, and cell adhesion (Fig. 2F). Additionally, GO analysis by cell compartment (CC) showed an enrichment of ‘lysosome’ and ‘extracellular space’ (Fig. S7B), further supporting the hypothesis that these cells are engulfing and digesting EVs. Notably, ∼30% of proteins with increased abundance in eCM-treated BMDMs were shared with those from the original efferocytic BMDM dataset of proteins with increased abundance, such as Adgre5, Clec4d, Edil3, Thbs1, ApoE and Fn1 (Fig. 2E and Fig. S8A). As expected, GO analysis of these shared proteins showed an enrichment of immune and inflammatory responses (Fig. S8B-C), highlighting the ability of efferocytic BMDMs to signal and cause phenotypic changes in neighbouring cells.

When further investigating these proteins with increased abundance, some of the most pronounced changes were from those associated with coagulation and complement, extracellular matrix, and platelet activation (Fig. 2G), indicative of a tissue remodelling phenotype. These processes have diverse yet intertwined functions, and importantly directly impact the innate immune system^19,20^. Platelet factor 4 (Pf4/Cxcl4), for example, is known to promote monocyte differentiation into macrophages^21^ and also decreases T cell proliferation and IFN-γ release in co-culture experiments^22^; here it was found to markedly increase in both efferocytic and eCM-treated BMDMs (Fig. 2G). Notably, it has previously been shown that activated platelets have an anti-inflammatory effect on macrophages^23^, demonstrative of the positive pro-resolutive feedback loop that may be induced by efferocytic macrophages. While it is a well-known marker of macrophages, CD11b is also a complement receptor that, in conjunction with FcγRs, plays a role in the clearance of necrotic cell debris^24^. Collectively, these data provide a novel insight to macrophage reprogramming events that occur after apoptotic cell clearance and how efferocytic macrophages can secrete factors that signal neighbouring cells.

## Discussion

Our study has been the first to successfully map global proteome and secretome changes in macrophages following efferocytosis. Using SILAC, we can effectively and efficiently separate the *Mus musculus* and *Homo sapiens* proteomes of macrophages and Jurkat cells, respectively. By combining this with DIA mass spectrometry, we have developed an easy-to-use assay that can provide deep proteome coverage. Importantly, our approach is compatible with various enrichment strategies to study post-translational modifications such as phosphorylation, glycosylation and ubiquitylation.

In support of our methodology, we identified several proteins that have previously been shown to play a positive role in efferocytosis, such as CD14^8^, thrombospondin 1^7^, Mfge8^10^, Slc7a11^11^, and Edil3/Del1^9^. Among the proteins found to increase in abundance, were proteins associated with M2 macrophage polarisation (Arg2) and macrophage recruitment (Adgre5/CD97), consistent with a shift towards a pro-resolving phenotype. Additionally, in support of the shift away from a pro-inflammatory phenotype, proteins found to have decreased abundance were pro-inflammatory molecules, such as MMP12, Clec12a, S100a9 and S100a6e. In our study, we have identified a plethora of putatively important proteins in processes of wound healing and inflammation, ideally, the resources provided here will be of great value to other researchers working in these fields. One such example is Notch2, whose abundance increased in efferocytic, as well as eCM-treated, BMDMs and has been shown to some degree to play a role in promoting diabetic wound healing when expressed on macrophages^25^.

The secretion of pro-resolving factors by both immune cells and apoptotic cells is an emerging theme in innate immunity^3,4^. Here, we applied our SILAC approach to study these secreted factors, in addition to their effect on naïve BMDMs. We also hypothesise that the secretome contains extracellular vesicles (EVs) that we speculate may contain both macrophage- and apoptotic cell-derived cargo. Future applications of our SILAC workflow will be used to elucidate the proteome of these EVs as well as those from apoptotic Jurkat cells only. While the efferocytic secretome is complex and contains proteins from both macrophage and apoptotic origin, we have shown that they are rich in pro-resolving molecules such as thrombospondin 1, Il1r2, and ApoE. Supportive of these findings is our observation that naïve BMDMs shift towards an efferocytic phenotype following 48 h incubation with efferocytosis conditioned medium. This unique phenotype is characterised by an increased abundance of proteins associated with extracellular matrix (ECM), complement and coagulation and platelet activation. While destruction of the ECM is detrimental to the 3D microenvironment, the dysregulation of ECM accumulation can lead to fibrosis. Therefore, the maintenance and fine-tuning of the ECM following inflammation is a crucial aspect of wound healing and the subsequent resolution of this inflammation^26^. Macrophage interactions with the ECM as well as cells such as fibroblasts are also an important aspect of this process^27^. It is possible that the increased abundance of specific proteins in eCM-treated BMDMs, such as Platelet factor 4/Cxcl4, a chemokine known to promote macrophage-fibroblast crosstalk, tissue remodelling and profibrotic phenotype in macrophages^28^, is driving them towards a tissue remodelling phenotype. Increased abundance of Fn1^28^, collagens and proteolgycans, and decreased abundance of MMP12^29,30^ in our datasets, further support this notion. Given the importance of balancing tissue-remodelling signals in order to prevent fibrosis, it is tempting to speculate that excessive efferocytosis, leading to an overactive feedback loop, may in fact contribute to a profibrotic phenotype. *In vivo* experiments whereby macrophages are overloaded with apoptotic cells and our SILAC workflow is applied, will help to further investigate this hypothesis.

Our data provides a comprehensive proteomic examination into the phenotype of BMDMs following the clearance apoptotic cells, as well as the proteins secreted by these cells that can exert an additional pro-resolutive effect on naïve cells, reminiscent of an alternative activation state. Our findings also reinforce the notion that the typical M1/M2 macrophage paradigm is relatively reductionist when describing the complex and dynamic environment that exists *in situ*. Notably, our novel and robust workflow presented here can be applied to *ex vivo* macrophages isolated from patients suffering from diseases where deficiencies in efferocytosis are known and will thus aid in a better understanding of the proteomic landscape of this important immunological process.

## Material and Methods

### Mice

Wild-type C57BL/6J mice were obtained from Charles River and bred under appropriate UK home office project licences.

### Cell culture

Bone marrow cells were collected from the tibia and femurs of male and female 6- to 8-week-old C57BL/6J wild-type mice. Cells were treated with red blood cell lysis buffer (155 mM NH_4_Cl, 12 mM NaHCO_3_, 0.1 mM EDTA) and plated on untreated 10 cm cell culture dishes (BD Biosciences) in IMDM (Gibco) containing 10% heat-inactivated FBS, 2 mM L-glutamine, 100 units/ml penicillin/streptomycin (Gibco) and 20% L929 conditioned media supplement. After 24 h, cells in the supernatant were transferred to untreated 10 cm Petri dishes (BD Biosciences) and left for 7 days, with additional media added every other day, for differentiation into bone marrow-derived macrophages (BMDMs)^31^. BMDMs were maintained in IMDM medium without L929 supernatant for at least 24 hours prior to the commencement of efferocytosis assays. BM A3.1A7 (BMA) immortalised macrophages were maintained in DMEM (Gibco) supplemented with 10% heat-inactivated FBS, 2mM L-glutamine and 100 units/ml penicillin/streptomycin (Gibco) and passaged every other day. Jurkat cells were maintained in RPMI (Gibco) supplemented with 10% heat-inactivated FBS, 2 mM L-glutamine and 100 units/ml penicillin/streptomycin (Gibco) and passaged every other day.

### Preparation and cell culture using SILAC

SILAC RPMI without L-Arginine and L-Lysine was supplemented with dialysed FBS (Gibco), 2 mM L-Glutamine, 100 units/ml Pen/Strep, 200 mg/L L-Proline (Sigma-Aldrich), and ‘heavy’-labelled L-Arginine (^13^C6; ^15^N4) at 84 mg/L and L-Lysine (^13^C6; ^15^N2) at 146 mg/L. The media was then filtered through a 0.2 μm membrane. Jurkat cells were grown in SILAC medium for a minimum of 8 days, passaging every other day, to allow incorporation of SILAC amino acids. A proportion of cells were taken for lysis in SDS lysis buffer, tryptic digestion, and data-independent (DIA) mass spectrometry (as described below), in order to assess the level of heavy label incorporation. Expressed as the number of peptides identified in each channel and the sum of intensities for each channel after channel Q-value filtering, ∼99% label incorporation was achieved (Fig. S1).

### Inducing apoptosis in Jurkat cells

On the day of the assay, Jurkat cells were collected by centrifugation at 500 xg for 5 min and resuspended in SILAC medium without supplementation and diluted to ∼1 - 5 x 10^6^ cells/ mL and treated with 1 μM Staurosporine (STS) or DMSO as a vehicle control. Cells were incubated for 1 or 3 h at 37°C. Jurkat cells were collected and washed 3x with serum-free media by centrifugation at 500 xg for 5 min to remove residual STS. Jurkat cells treated with 1 μM STS (or vehicle control) for 3 h were resuspended in warm normal IMDM for efferocytosis assays, and those treated for 1 or 3 h were taken for flow cytometry analysis as described below.

### Live/Dead assay using flow cytometry

Cell viability of Jurkat cells following STS or DMSO treatment was tested using BD Pharmingen™ PE Annexin V Apoptosis Detection Kit I with no modifications to the protocol. Briefly, STS- or DMSO-treated Jurkat cells were resuspended in 1x Binding Buffer at a concentration of 1 x 10^6^ cells/ml. PE Annexin V and 7-AAD was added, tubes vortexed, and incubated for 15 min at RT in the dark. Samples were immediately analysed on a BD FACSymphony™ A1 Cell Analyzer. Data was analysed using FlowJo software to perform compensation and gate on single double positive events after excluding doublets.

### Efferocytosis assay for proteomics

Macrophages were seeded at ∼1.0 x 10^6^ cells/ well in 6-well plates and allowed to adhere overnight. STS- or DMSO-treated Jurkat cells were diluted to appropriate densities for an MOI of 4:1. Macrophage media was aspirated and replaced with fresh warm medium containing Jurkat cells. Plates were centrifuged at 200 xg for 1 min and immediately placed at 37°C for 30 min to allow attachment/engulfment (“pulse”). After 30 min, media was aspirated, and cells washed gently 2x with warm PBS and replaced with fresh warm IMDM. Plates were incubated for a further 30 min or 18 h at 37°C (“chase”). After each time point, cells were washed twice with ice cold PBS and lysed in 5% SDS/ 50 mM TEAB, pH 7.55 lysis buffer containing cOmplete protease inhibitors (Roche) and universal nuclease (Pierce) with scraping.

### Collection of supernatants for secretome analysis

Efferocytosis assays were performed as described as above with a few modifications. One hour prior to the assay, BMDM medium was replaced with serum-free IMDM to equilibrate them, and all subsequent steps were performed in serum-free IMDM. Additionally, controls did not receive any apoptotic Jurkat cells but were otherwise treated exactly the same as efferocytosis conditions. Supernatants were collected 6 h and 18 h following the ‘chase’ step. Once collected, supernatants were centrifuged at 1,000 xg for 10 minutes to remove cells and debris. Proteins were then precipitated using the methanol/chloroform method. Briefly, methanol was added to supernatants, followed by chloroform and vortexing to mix. H_2_O was added, samples were vortexed and centrifuged at 4,000 xg for 15 min at 4°C. The upper layer was discarded, and methanol added to the sample (bottom phase plus protein interphase containing protein). Samples were centrifuged at 14,000 xg for 30 min at 4°C and the supernatant discarded. Pellets were resuspended in lysis buffer with heating at 70°C and vigorous vortexing for 20 min.

### Treatment of BMDMs with efferocytosis conditioned medium

Untreated BMDMs were seeded into 6-well plates as described above and left to adhere overnight. Prior to the addition of conditioned medium (CM), these cells were washed once in PBS to remove any non-adherent cells. CM was collected from identical BMDMs 18 h after efferocytosis (efferocytosis CM) or no treatment (control CM), centrifuged at 1,000 xg for 10 minutes, and directly added to plates containing BMDMs. The only modification to the efferocytosis assays in this case, was that they were performed in the presence of 10% FCS. BMDMs were incubated with control or efferocytosis CM for 48 h, after which they were washed with PBS and lysed directly in 5% SDS/ 50 mM TEAB, pH 7.55 containing cOmplete protease inhibitors (Roche) and universal nuclease (Pierce).

### Sample preparation for mass spectrometry analysis

Protein lysates were centrifuged at 16,000 xg using a benchtop centrifuge for 20 min to remove any insoluble material. The supernatant was collected, and protein concentrations quantified using Pierce BCA Protein Assay Kit (Thermo Fisher Scientific). Samples were reduced with 10 mM tris(2-carboxyethyl)phosphine (TCEP) for 20 min at 37°C, and then alkylated with 10 mM iodoacetamide (Sigma) for 20 min in the dark. Samples were then digested with trypsin using the S-trap protocol (ProtiFi) as previously described^31^. Resulting peptides were dried down using a SpeedVac concentrator (Thermo Scientific).

### Mass spectrometry analysis

Dried down peptide samples were resuspended in HPLC-grade water containing 0.1% formic acid at a concentration of 500 ng/μl and placed into glass autosampler vials. Liquid chromatography (LC) was performed using a NanoElute II (Bruker Daltonics) with a 15 cm Aurora Elite C18 column with integrated captive spray emitter (IonOpticks), at 50 °C, and a pre-column 5 mM PepMapTM C18 trap (ThermoFisher Scientific). Buffer A was 0.1 % formic acid in HPLC water, buffer B was 0.1 % formic acid in acetonitrile. Immediately prior to LC-MS, peptides were resuspended in buffer A and a volume of peptides equivalent to 500 ng was injected into the system, and subjected to a gradient of 5-35 % buffer B, 300 nl/min, for 60 minutes, followed by increasing buffer B to 95 % in 30 seconds, maintaining buffer B at 95 % for 4.5 min, and increasing the flow rate to 600 nl/min during this time. Equilibration with buffer A was performed with 10 and 4 column volumes for the trap and separation column, respectively. The LC was used in line with a timsToF-HT mass spectrometer (Bruker Daltonics) operated in DIA-PASEF mode. Mass and IM ranges for PASEF were 300-1200 m/z and 0.6-1.4 1/K0, DIA-PASEF was performed using variable width IM-m/z windows, designed using py_diAID ^32^, and are provided (see data sharing section). TIMS ramp and accumulation times were 100 ms each with a Ramp rate of 9.42 Hz, total cycle time was ∼1.8 seconds. Collision energy was applied in a linear fashion, where ion mobility = 0.6-1.6 1/K0, and collision energy = 20 - 59 eV.

### Data analysis

Raw data files were initially searched using DIA-NN 1.8.1 to generate an *in silico* spectral library, necessary for searching SILAC samples, from the *Mus musculus* (downloaded from Uniprot on the 21^st^ of July 2023 containing 17,167 entries) and *Homo sapiens* (downloaded from Uniprot on the 15^th^ of August 2023, containing 20,423 entries) proteomes, as well as a common contaminants list. This library was generated in DIA-NN 1.8.1 with fixed carbamidomethyl (C) and variable oxidation (M) peptide mass modifications, and mass accuracy and MS1 accuracy set to 10. Peptide length range was 7 – 30, precursor charge range 1 – 4, precursor m/z range 300 – 1800, and fragment ion m/z range 200 – 1800. For our SILAC searches using the spectral libraries generated above, the same parameters were used in addition to the following modifications added to the search window: --fixed-mod SILAC,0.0,KR,label, --lib-fixed-mod SILAC, --channels SILAC,L,KR,0:0; SILAC,H,KR,8.014199:10.008269, --peak-translation, --original-mods, --matrix-ch-qvalue and --matrix-tr-qvalue. Report files were then processed using RStudio to separate channels, followed by protein quantification using the iq package^33^ to calculate the MaxLFQ based on Precursor Translated intensities, with the Channel q-value threshold set to 0.01. Following the removal of contaminants, proteins only identified by one unique peptide and any proteins not found in at least 3 (or 4 in the case of supernatant and conditioned medium experiments) replicates of one group, the data was normalised based on median intensities, log2 transformed, and fold-changes calculated with the limma package using the BH p-value adjust method^34^.

### Efferocytosis assay for flow cytometry

Efferocytosis assays were performed as described above, with the following modifications. BMDMs, seeded into 6-well plates, and apoptotic Jurkat cells were first stained with CellTrace Violet (Thermo Fisher Scientific) and CellTracker Deep Red (Thermo Fisher Scientific), respectively. Following the 30 min “pulse”, efferocytosis was allowed to occur for 30 min or 3 h before cells were washed with PBS and detached using trypsin (Fig. 1A). Cells were then washed with PBS by centrifugation and fixed using 4% paraformaldehyde for 15 min at room temperature. Cells were washed with PBS by centrifugation, resuspended in PBS containing 2% FCS (FACS buffer), and run on a FACSymphony A5 Cell Analyzer (BD Biosciences) using the 405 nm laser with 450/50 bandpass filter, and 635 nm laser with 670/30 bandpass filter. Data was analysed using FlowJo software to gate on double positive events after excluding doublets.

### Efferocytosis assay for confocal microscopy

Efferocytosis assays were performed as described for flow cytometry with the following modifications. Prior to performing the assay, BMDMs were seeded onto No. 1.5H coverslips (Marienfeld-Superior) and allowed to adhere overnight. Following efferocytosis, cells were washed with PBS and fixed using 4% paraformaldehyde for 15 min at room temperature. Cells were washed and mounted onto slides using ProLong Glass (Thermo Fisher Scientific). Samples were imaged using a Zeiss LSM 800 at 63x magnification in confocal mode using lasers 405 nm and 647 nm, both set at 1% laser power with 650V detector gain. Bidirectional scan direction was used with 8x averaging, imaging at 16-bit depth. Image analysis was performed using Imaris software.

### Purification of extracellular vesicles using ultracentrifugation

Efferocytosis assays were performed as described above for total cell proteomics experiments. Following 18 h, supernatants from each well of a 6-well plate per replicate (for both control and efferocytosis experiments) were collected and combined. Samples were kept on ice at all times, and all subsequent centrifugation steps were also performed at 4°C. Samples were then sequentially centrifuged at i) 400 xg for 5 min to remove cells, ii) 2,000 xg for 20 min to remove cell debris, and iii) 10,000 xg for 45 min to remove apoptotic bodies. Samples were then centrifuged at 100,000 xg for 90 min to pellet EVs. Supernatants were discarded, and EVs were washed by centrifugation at 100,000 xg for 90 min. Pellets were resuspended in PBS and stored at −80°C.

### Transmission electron microscopy of extracellular vesicles

Ten microlitres of EVs in solution were placed on a carbon-coated, glow discharged copper grid (Gilder Grids) for 1 min. Subsequently, the excess from the droplet was wicked away by touching to the edge of a hardened filter paper. The sample was negatively stained by dropping the grid, sample side down, onto a droplet of 2% aqueous uranyl acetate (Agar Scientific) for a few seconds. Grid was dried by filter paper again and air dried under a heat lamp prior to viewing. Grids were examined on a Hitachi HT7800 transmission electron microscope using an Emsis Xarosa camera with Radius software.

### Statistical analysis

Statistical analyses were performed using RStudio built on R version 4.3.0 and GraphPad 9.5.0. For Volcano plots created using GraphPad, adjusted p-value cut-offs were set at *P* < 0.05 (-log10 value of 1.3) and fold-change cut-offs set at +/- 1.5 (log2 value of +/- 0.5849). Further analysis was performed in RStudio to produce heatmaps (pheatmap) and circle plots of gene ontology (GO) enrichments (GOplot) performed using David^35,36^.

### Data availability

The mass spectrometry proteomics data have been deposited to the ProteomeXchange Consortium^37^ via the PRIDE^38^ partner repository with the dataset identifier PXD047180.

## Author contributions

BBAR and MT conceptualised the study and wrote the manuscript. BBAR performed and analysed most experiments. AMF performed some experiments and assisted with data analysis.

## Acknowledgements

We would like to thank the Newcastle University Comparative Biology Centre, Flow Cytometry Core Facility, Electron Microscopy Research Services (specific thanks to Tracey Davey), and the BioImaging Unit (specific thanks to David Bulmer) for their support and assistance in this work. MT, BBAR, and AMF are funded by a Wellcome Trust Investigator Award to MT (215542/Z/19/Z). This study was also partly funded by a Research Excellence Research Award from Newcastle University to BBAR (NU-017983). The Hitachi HT7800 transmission electron microscope used in this study was funded by the Biotechnology and Biological Sciences Research Council (grant reference BB/R013942/1). The timsTOF HT mass spectrometer used in this study was part-funded by the Wellcome Trust (212947/Z/18/Z).

