## Supplementary Figures and Tables for "Proteomic mapping of macrophages in response to the clearance of apoptotic cells reveals a unique reprogramming profile"

### Supplementary figures & legends

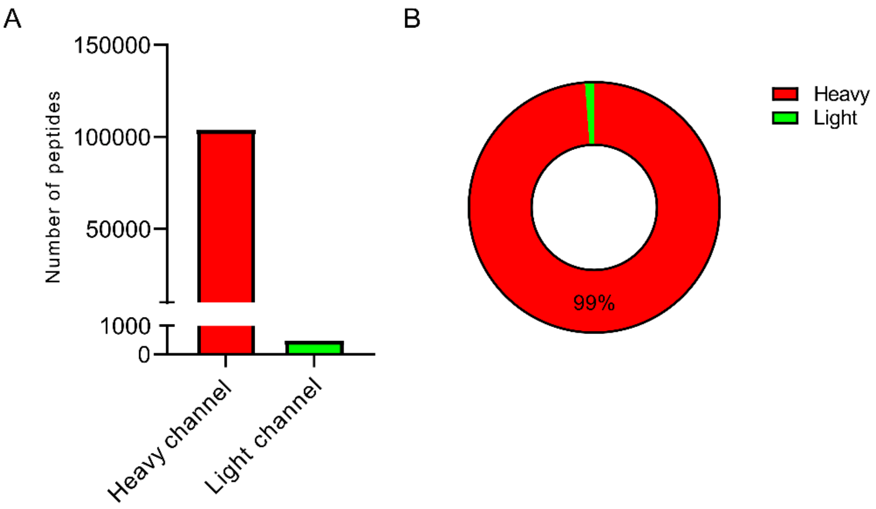


**Supplementary Figure 1:** Mass spectrometry validation of SILAC amino acid incorporation by Jurkat cells after 8 days of culture. (A) Number of 'heavy' or 'light' labelled peptides identified after applying 1% channel Q-value cut-offs. (B) Sum of intensities of 'heavy' or 'light' labelled peptides after applying 1% channel Q-value cut-offs, demonstrating that ~99% of peptides were SILAC-labelled.

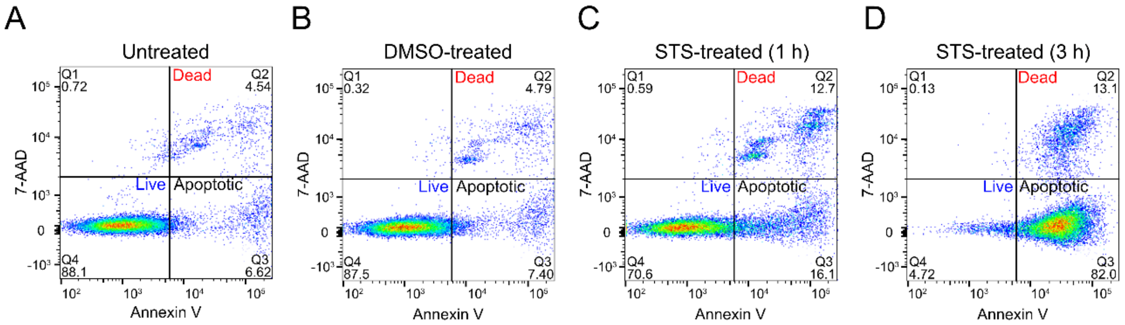


**Supplementary Figure 2:** Flow cytometry demonstrating that staurosporine (STS) treatment induces early apoptosis after 3 h (D), while little apoptosis was observed in untreated (A), vehicle control (B) and 1 h (C) STS treatment conditions.

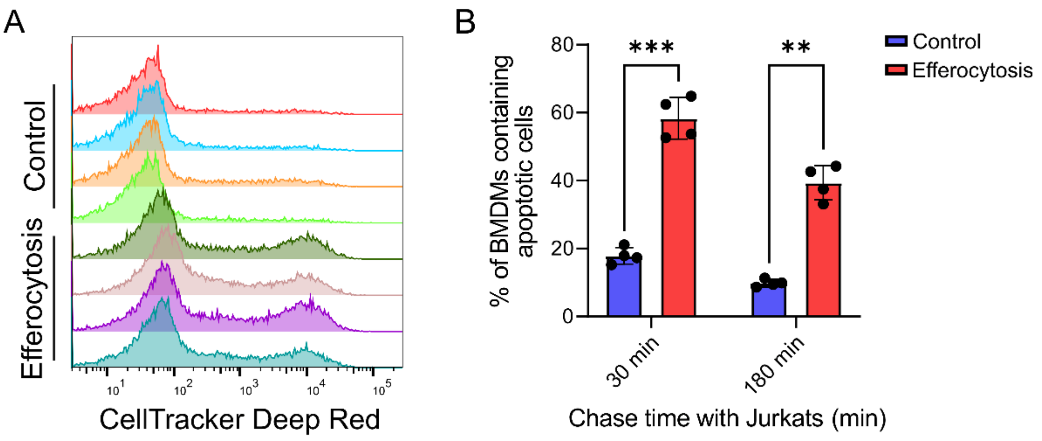


**Supplementary Figure 3:** Flow cytometry assay demonstrating the uptake of apoptotic cells by macrophages after 30 min and 3 h. (A) Histogram showing CellTracker Deep Red fluorescence intensity after gating for CellTracker Violet positive events in control (BMDMs + DMSO-treated Jurkat cells) and efferocytosis (BMDMs + STS-treated Jurkat cells) conditions after 30 min. (B) Quantification of CellTracker Violet/Deep Red double positive events, indicative of efferocytosis, after 30 min and 3 h. Data is presented as the average of 4 biological replicates with standard deviation. ***P*-value $< 0.01$; ****P*-value $< 0.001$ by paired two-tailed Student's *t*-test.

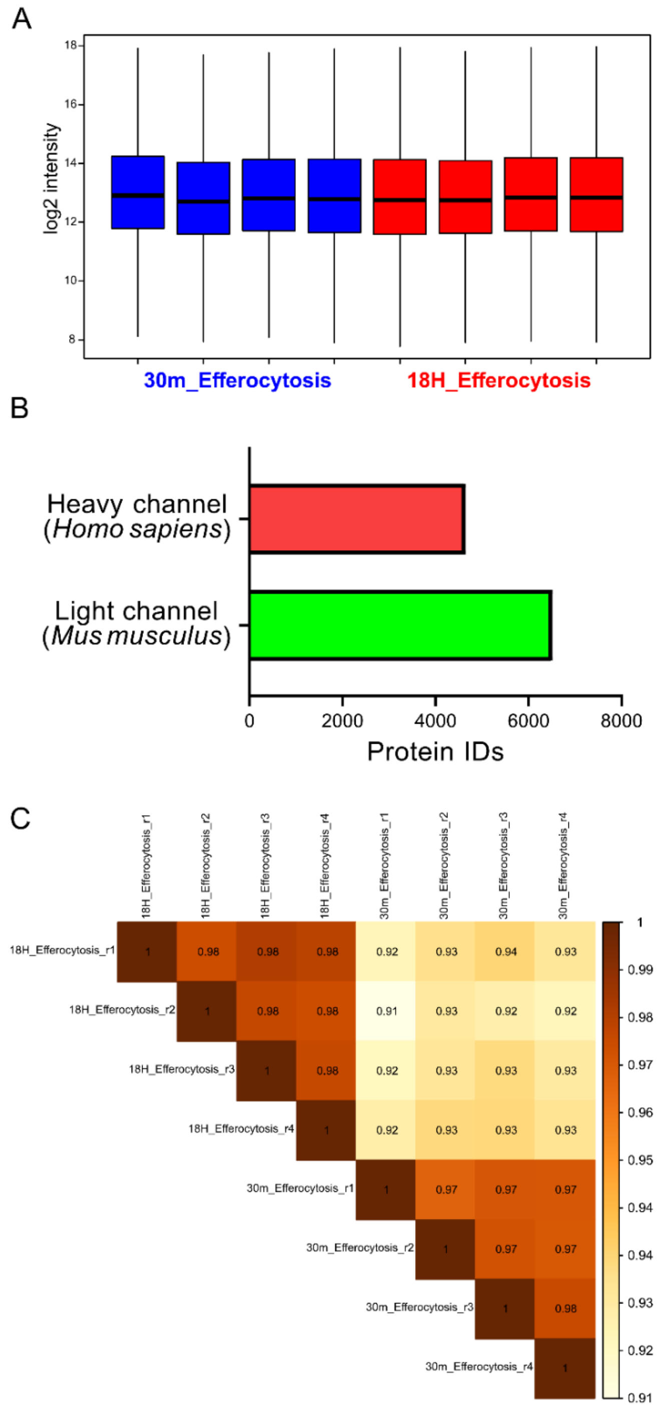


**Supplementary Figure 4:** Quality control of mass spectrometry data from total cell efferocytosis experiments. (A) Box and whisker plot of non-normalised, log2 transformed, intensities from the light channel of *Mus musculus* proteins from 4 biological replicates, showing that there is little inter-sample variability. Whiskers represent the minimum and maximum values, and central bands indicate the median. (B) Bar graph showing the number of protein identifications, post-filtering, belonging to *Mus*

*musculus* and *Homo sapiens* from the ‘light’ and ‘heavy’ channels, respectively. (C) Correlation plots between control (30 m) and efferocytosis (18 h) conditions demonstrating that there is little inter-sample variability. Correlation coefficients are shown in each box and the colour scale is on the right.

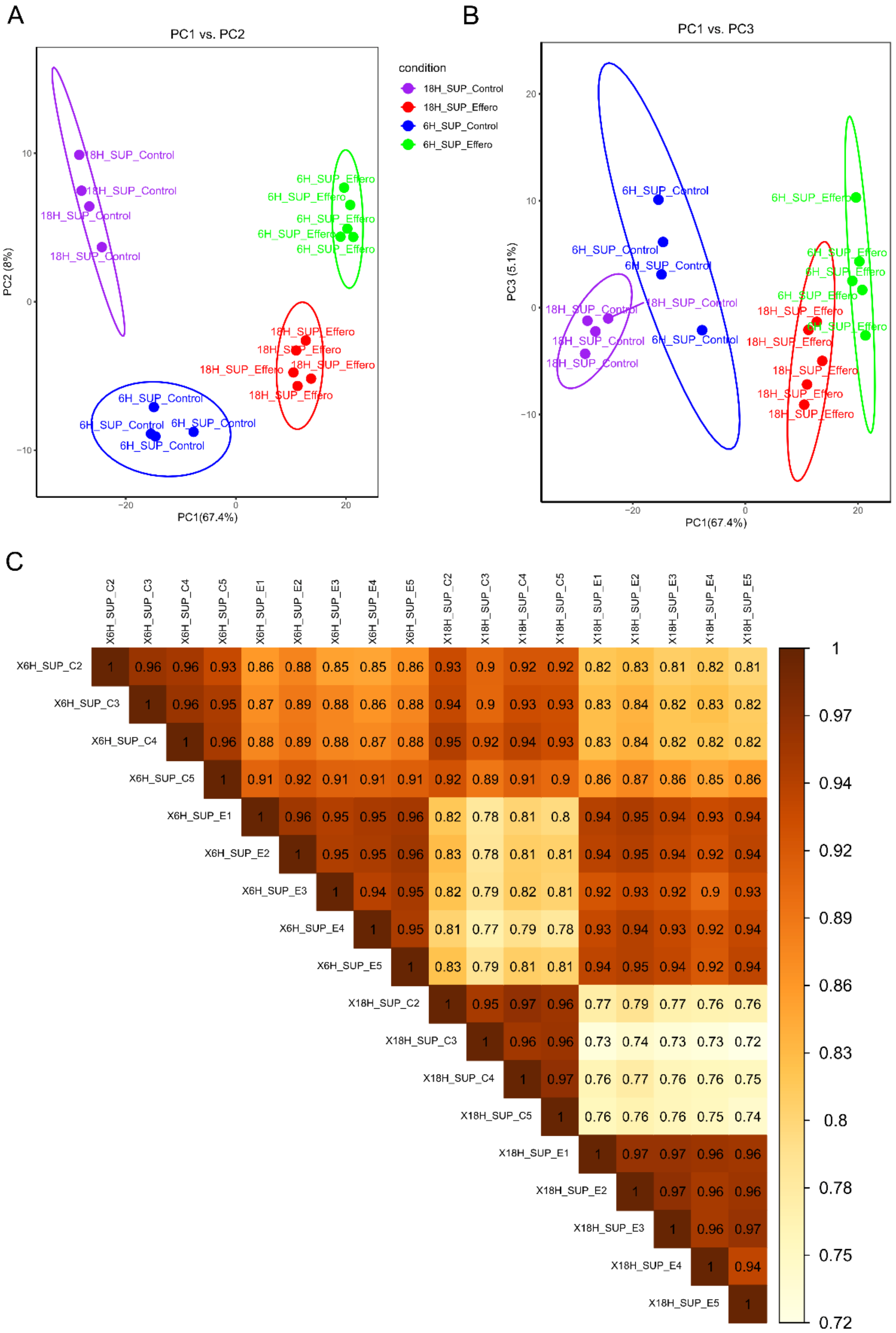


**Supplementary Figure 5:** Quality control of mass spectrometry data from secretome experiments. Principal component analyses (A-B) and correlation plots (C) of the BMDM secretomes across all experimental conditions. (A) PC1 vs PC2. (B) PC1 vs PC3. (C) Correlation coefficients are show in each box and the colour scale is on the right.

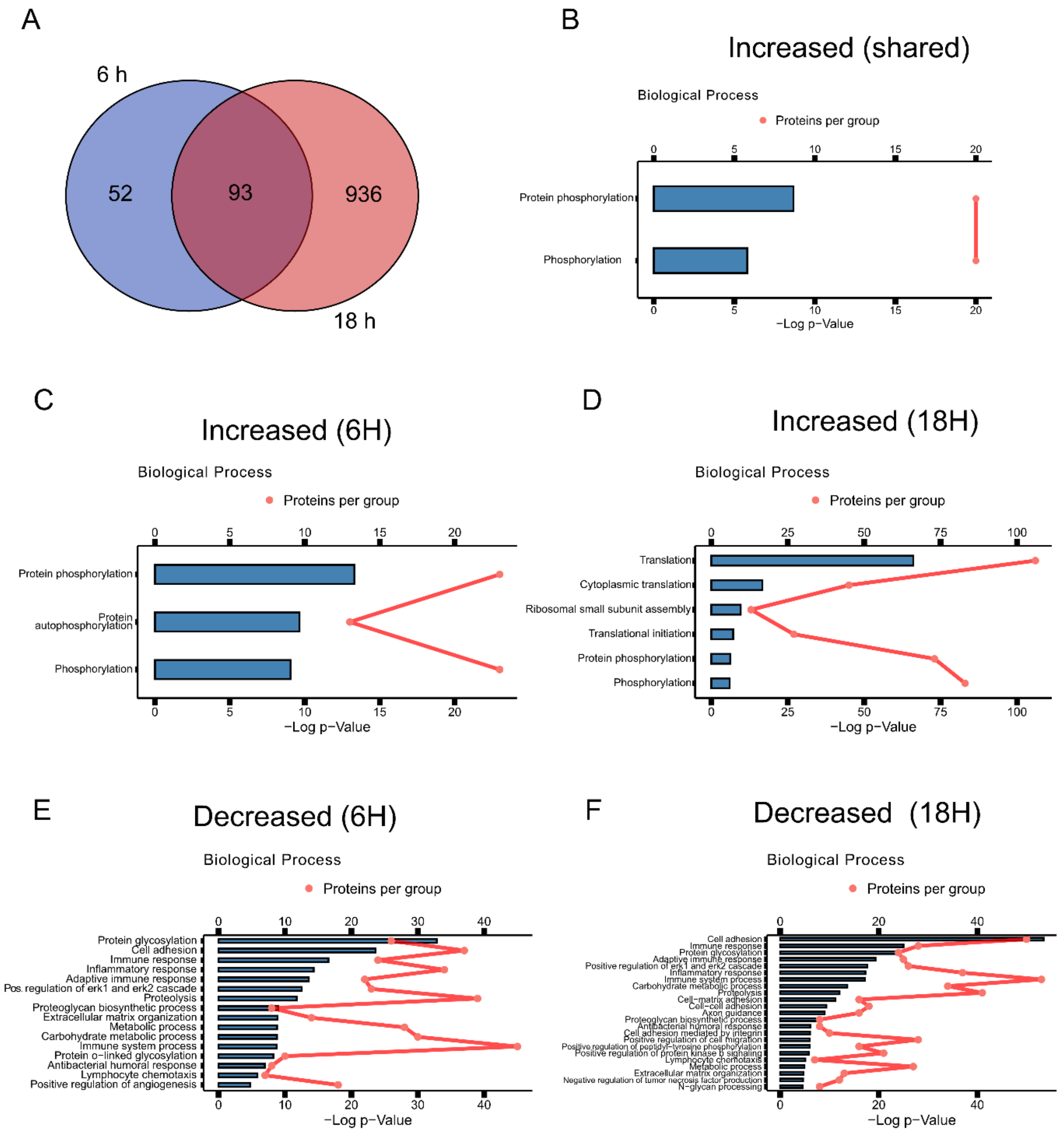


**Supplementary Figure 6:** Analysis of BMDM proteins from secretomes with increased abundance following efferocytosis. (A) Venn diagram demonstrating the overlap of proteins with increased abundances from supernatants following 6 and 18 h efferocytosis. (B-F) Gene ontology enrichment by biological process for: shared increased (B), increased at 6 h (C), increased at 18 h (D), decreased at 6 h (E), decreased at 18 h (F).

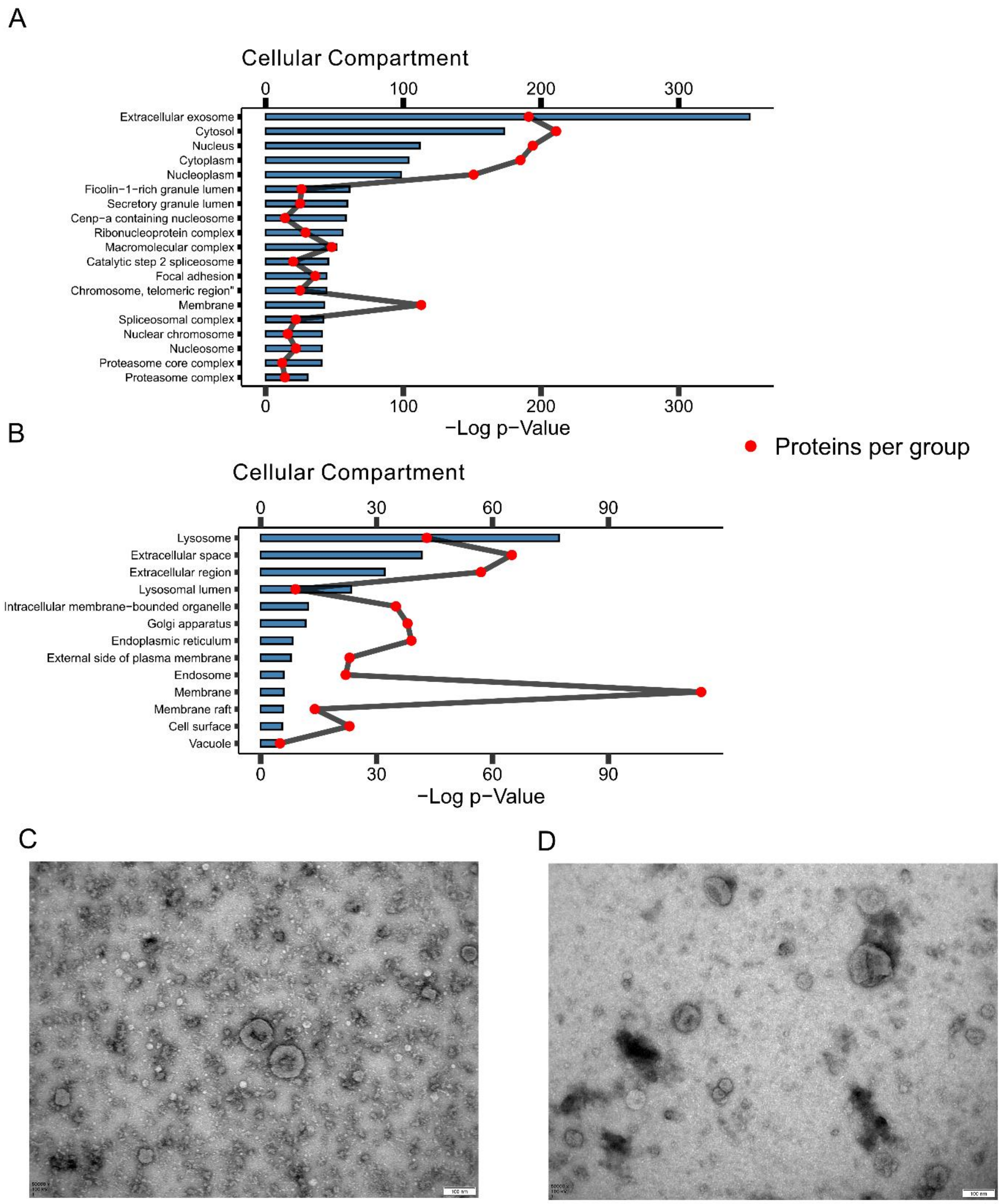


**Supplementary Figure 7:** Secretion of extracellular vesicles by BMDMs following efferocytosis. (A) Gene ontology analysis of heavy-labelled human proteins from 18 h secretomes by cellular compartment, showing a strong enrichment of proteins belonging to 'extracellular exosome'. (B) Gene ontology analysis of light-labelled murine proteins from eCM-treated BMDMs by cellular component showing a strong enrichment of lysosome. (C-D) Transmission electron microscopy of extracellular vesicle (EVs) preparations from 18 h control and efferocytosis supernatants. EVs were prepared by ultracentrifugation after first removing cells, cell debris, and apoptotic bodies by sequential centrifugation. Scale bars are 100 μm.

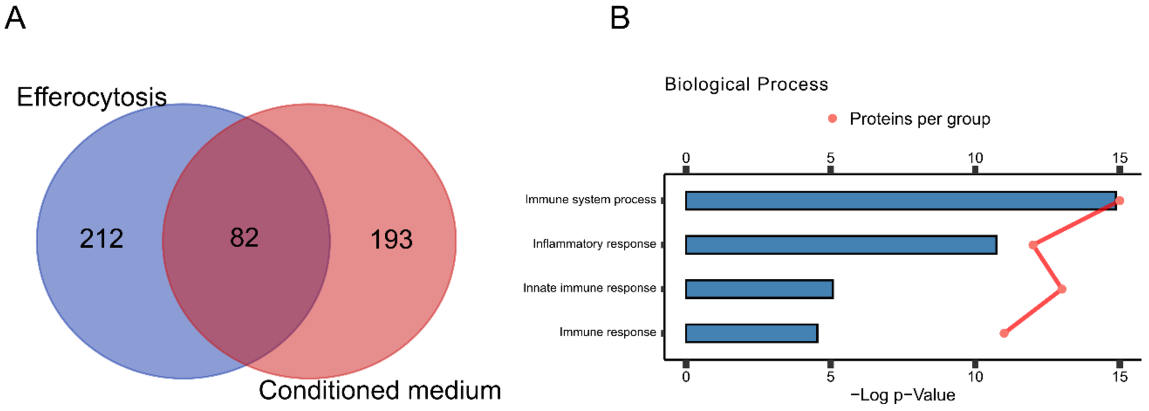


C

| Uniprot ID | Gene name | Protein name |
|---|---|---|
| A2A690 | TANC2 | Protein TANC2 |
| A8Y5H7 | S14L1 | SEC14-like protein 1 |
| B8JK39 | ITA9 | Integrin alpha-9 |
| E9Q414 | APOB | Apolipoprotein B-100 |
| E9Q612 | PTPRO | Receptor-type tyrosine-protein phosphatase O |
| O35474 | EDIL3 | EGF-like repeat and discoidin I-like domain-containing protein 3 |
| O35516 | NOTC2 | Neurogenic locus notch homolog protein 2 |
| O54918 | B2L11 | Bcl-2-like protein 11 |
| O88668 | CREG1 | Protein CREG1 |
| O88935 | SYN1 | Synapsin-1 |
| P04186 | CFAB | Complement factor B |
| P08226 | APOE | Apolipoprotein E |
| P09581 | CSF1R | Macrophage colony-stimulating factor 1 receptor |
| P10810 | CD14 | Monocyte differentiation antigen CD14 |
| P11276 | FINC | Fibronectin |
| P11680 | PROP | Properdin |
| P15105 | GLNA | Glutamine synthetase |
| P15261 | INGR1 | Interferon gamma receptor 1 |
| P21447 | MDR1A | ATP-dependent translocase ABCB1 |
| P21956 | MFGM | Lactadherin |
| P21981 | TGM2 | Protein-glutamine gamma-glutamyltransferase 2 |
| P21995 | EMB | Embigin |
| P25119 | TNR1B | Tumor necrosis factor receptor superfamily member 1B |
| P25446 | TNR6 | Tumor necrosis factor receptor superfamily member 6 |
| P28033 | CEBPB | CCAAT/enhancer-binding protein beta |
| P32261 | ANT3 | Antithrombin-III |
| P35441 | TSP1 | Thrombospondin-1 |
| P41233 | ABCA1 | Phospholipid-transporting ATPase ABCA1 |
| P42082 | CD86 | T-lymphocyte activation antigen CD86 |
| P43883 | PLIN2 | Perilipin-2 |
| P45377 | ALD2 | Aldose reductase-related protein 2 |
| P50543 | S10AB | Protein S100-A11 |
| P51670 | CCL9 | C-C motif chemokine 9 |
| P55065 | PLTP | Phospholipid transfer protein |
| P57787 | MOT4 | Monocarboxylate transporter 4 |
| P58242 | ASM3B | Acid sphingomyelinase-like phosphodiesterase 3b |
| P62746 | RHOB | Rho-related GTP-binding protein RhoB |
| P99029 | PRDX5 | Peroxiredoxin-5, mitochondrial |
| Q08093 | CNN2 | Calponin-2 |
| Q09014 | NCF1 | Neutrophil cytosol factor 1 |
| Q3TNL8 | IPRI | Inositol 1,4,5-trisphosphate receptor-interacting protein |

| Uniprot ID | Gene name | Protein name |
|---|---|---|
| Q3TWN3 | CNNM2 | Metal transporter CNNM2 |
| Q3URF8 | KCD21 | BTB/POZ domain-containing protein KCTD21 |
| Q60780 | GAS7 | Growth arrest-specific protein 7 |
| Q60963 | PAFA | Platelet-activating factor acetylhydrolase |
| Q61333 | TNAP2 | Tumor necrosis factor alpha-induced protein 2 |
| Q61409 | PDE3B | cGMP-inhibited 3',5'-cyclic phosphodiesterase 3B |
| Q61508 | ECM1 | Extracellular matrix protein 1 |
| Q61703 | ITIH2 | Inter-alpha-trypsin inhibitor heavy chain H2 |
| Q61838 | PZP | Pregnancy zone protein |
| Q61878 | PRG2 | Bone marrow proteoglycan |
| Q62431 | ARI3A | AT-rich interactive domain-containing protein 3A |
| Q6DFX2 | ANTR2 | Anthrax toxin receptor 2 |
| Q8BG48 | ST17B | Serine/threonine-protein kinase 17B |
| Q8BGR2 | LRC8D | Volume-regulated anion channel subunit LRRC8D |
| Q8BJQ4 | TCAM2 | TIR domain-containing adapter molecule 2 |
| Q8BQQ1 | ZDH14 | Palmitoyltransferase ZDHHC14 |
| Q8BQS4 | EEIG2 | EEIG family member 2 |
| Q8BW49 | TTC12 | Tetratricopeptide repeat protein 12 |
| Q8BYY4 | TT39B | Tetratricopeptide repeat protein 39B |
| Q8C0T5 | SI1L1 | Signal-induced proliferation-associated 1-like protein 1 |
| Q8CE08 | PPAP | Prostatic acid phosphatase |
| Q8CJC5 | NEUL3 | E3 ubiquitin-protein ligase NEURL3 |
| Q8K1H7 | T11L2 | T-complex protein 11-like protein 2 |
| Q8K3J9 | GPC5C | G-protein coupled receptor family C group 5 member C |
| Q8K482 | EMIL2 | EMILIN-2 |
| Q921I1 | TRFE | Serotransferrin |
| Q99LE1 | RIPL2 | RILP-like protein 2 |
| Q99M71 | EPDR1 | Mammalian ependymin-related protein 1 |
| Q9CRB6 | TPPP3 | Tubulin polymerization-promoting protein family member 3 |
| Q9DBS1 | TMM43 | Transmembrane protein 43 |
| Q9EQC1 | 3BHS7 | 3 beta-hydroxysteroid dehydrogenase type 7 |
| Q9JJZ6 | KLF13 | Krueppel-like factor 13 |
| Q9JKF4 | CLC6A | C-type lectin domain family 6 member A |
| Q9JL95 | PRG3 | Proteoglycan 3 |
| Q9JLC3 | MSRB1 | Methionine-R-sulfoxide reductase B1 |
| Q9QUN7 | TLR2 | Toll-like receptor 2 |
| Q9QX11 | CYH1 | Cytohesin-1 |
| Q9WVK4 | EHD1 | EH domain-containing protein 1 |
| Q9Z0M6 | AGRE5 | Adhesion G protein-coupled receptor E5 |
| Q9Z126 | PLF4 | Platelet factor 4 |
| Q9Z2H6 | CLC4D | C-type lectin domain family 4 member D |

**Supplementary Figure 8:** eCM-treated BMDMs resemble efferocytic BMDMs. (A) Venn diagram of shared proteins with increased abundance between efferocytosis (total cells) and eCM-treated BMDMs.

(B) Gene ontology analysis of shared proteins by biological process showing an enrichment of immune system processes. (C) Table of shared proteins, with Uniprot IDs and gene/protein names listed.

**Supplementary Table 1: Proteins present/absent from 3 out of 4 replicates of one condition from efferocytosis experiments. MaxLFQ calculated intensities are reported for each replicate.**

| Protein.Group | Genes | 30m_ r1 | 30m_ r2 | 30m_ r3 | 30m_ r4 | 18H_ r1 | 18H_ r2 | 18H_ r3 | 18H_ r4 |
|---|---|---|---|---|---|---|---|---|---|
| Q8BM75 | Arid5b | 10.111 | 10.4975 | 10.3108 | 10.4177 | 0 | 0 | 0 | 0 |
| Q9DAP7 | Asf1b | 11.0989 | 10.8958 | 10.8636 | 10.7952 | 0 | 0 | 0 | 0 |
| P97477 | Aurka | 9.98999 | 10.0721 | 10.7631 | 10.663 | 0 | 0 | 0 | 0 |
| Q14AI0 | DSCC1 | 10.013 | 9.89207 | 10.3801 | 10.3417 | 0 | 0 | 0 | 0 |
| Q8C2P3 | Dus1l | 10.4013 | 0 | 9.9978 | 10.3929 | 0 | 0 | 0 | 0 |
| P04231 | H2-Eb1 | 12.2903 | 12.0967 | 12.4606 | 12.4785 | 0 | 0 | 0 | 0 |
| Q60848 | Hells | 11.1489 | 11.5766 | 11.8614 | 11.6017 | 0 | 0 | 0 | 0 |
| Q8R1M0 | Hmces | 10.547 | 9.41668 | 11.6501 | 0 | 0 | 0 | 0 | 0 |
| Q6IE82 | Jade3 | 11.4237 | 11.7424 | 0 | 11.3419 | 0 | 0 | 0 | 0 |
| Q80WQ8 | Mis18bp1 | 14.3272 | 0 | 14.3409 | 14.2624 | 0 | 0 | 0 | 0 |
| Q6GQX2 | Nckap5l | 11.2677 | 10.9357 | 10.8646 | 11.2008 | 0 | 0 | 0 | 0 |
| P51954 | Nek1 | 11.7435 | 11.7033 | 11.6802 | 0 | 0 | 0 | 0 | 0 |
| Q9ERH4 | Nusap1 | 11.0068 | 10.6252 | 10.5578 | 10.4463 | 0 | 0 | 0 | 0 |
| Q9JJ78 | Pbk | 11.8443 | 12.2244 | 10.8982 | 12.3502 | 0 | 0 | 0 | 0 |
| Q9CQX4 | Pclaf | 9.77517 | 9.46618 | 9.60661 | 9.67408 | 0 | 0 | 0 | 0 |
| Q9WVF7 | Pole | 11.5603 | 10.6776 | 9.45597 | 9.63715 | 0 | 0 | 0 | 0 |
| Q9WVM1 | Racgap1 | 10.5126 | 0 | 11.1081 | 10.7879 | 0 | 0 | 0 | 0 |
| Q08297 | Rad51 | 10.712 | 9.93593 | 9.38241 | 11.2971 | 0 | 0 | 0 | 0 |
| Q60695 | Rgl1 | 11.7461 | 11.4381 | 11.6366 | 11.7066 | 0 | 0 | 0 | 0 |
| Q9ER80 | Rtp4 | 10.7058 | 10.5736 | 0 | 10.6634 | 0 | 0 | 0 | 0 |
| P97440 | Slbp | 12.5586 | 10.6481 | 9.87437 | 10.5407 | 0 | 0 | 0 | 0 |
| Q9R1X4 | Timeless | 12.4227 | 11.4559 | 0 | 10.9934 | 0 | 0 | 0 | 0 |
| P04184 | Tk1 | 11.1854 | 11.073 | 11.0696 | 11.0121 | 0 | 0 | 0 | 0 |
| Q6ZQF0 | Topbp1 | 9.43884 | 0 | 10.3471 | 9.04293 | 0 | 0 | 0 | 0 |
| A2APB8 | Tpx2 | 9.51145 | 10.5762 | 9.41768 | 10.2207 | 0 | 0 | 0 | 0 |
| Q8BXJ2 | Trerf1 | 10.9793 | 11.3926 | 11.3029 | 10.7518 | 0 | 0 | 0 | 0 |
| P07607 | Tyms | 11.865 | 12.1238 | 12.6747 | 11.9997 | 0 | 0 | 0 | 0 |
| Q9QZU9 | Ube2l6 | 13.135 | 0 | 13.1742 | 13.1682 | 0 | 0 | 0 | 0 |
| Q8VDF2 | Uhrf1 | 11.7901 | 11.6353 | 11.3466 | 11.3016 | 0 | 0 | 0 | 0 |
| B1AQJ2 | Usp36 | 0 | 9.44716 | 10.8608 | 10.5265 | 0 | 0 | 0 | 0 |
| A6PWY4 | Wdr76 | 11.8119 | 12.2809 | 12.5001 | 11.9921 | 0 | 0 | 0 | 0 |
| Q6NZP1 | Zranb3 | 9.5647 | 9.8688 | 9.5465 | 0 | 0 | 0 | 0 | 0 |
| P53657 | Pklr | 0 | 0 | 0 | 0 | 0 | 11.4102 | 11.1252 | 11.5259 |
| Q7TMM8 | Parp16 | 0 | 0 | 0 | 0 | 0 | 11.9838 | 10.9489 | 11.1856 |
| Q6NS86 | Znf366 | 0 | 0 | 0 | 0 | 12.2201 | 12.9681 | 12.5735 | 12.4637 |

| | | | | | | | | | |
|---|---|---|---|---|---|---|---|---|---|
| O88968 | Tcn2 | 0 | 0 | 0 | 0 | 11.6308 | 11.6439 | 11.7898 | 11.1018 |
| P11627 | L1cam | 0 | 0 | 0 | 0 | 11.4986 | 11.4652 | 11.8576 | 11.5235 |
| Q61493 | Rev3l | 0 | 0 | 0 | 0 | 11.3904 | 11.3128 | 11.5447 | 0 |
| Q61503 | Nt5e | 0 | 0 | 0 | 0 | 11.273 | 11.1814 | 11.1911 | 11.0666 |
| P42228 | Stat4 | 0 | 0 | 0 | 0 | 11.1624 | 10.9734 | 11.0742 | 11.2233 |
| Q3UQ28 | Pxdn | 0 | 0 | 0 | 0 | 11.0951 | 11.6395 | 10.7895 | 11.0774 |
| P35917 | Flt4 | 0 | 0 | 0 | 0 | 11.0612 | 11.165 | 10.4149 | 11.1206 |
| P33766 | Fpr1 | 0 | 0 | 0 | 0 | 10.9767 | 0 | 11.6612 | 12.0765 |
| A2A8L5 | Ptprf | 0 | 0 | 0 | 0 | 10.9375 | 11.3581 | 11.1785 | 11.2976 |
| Q9Z0P7 | Sufu | 0 | 0 | 0 | 0 | 10.8489 | 10.6527 | 10.7803 | 10.595 |
| P27784 | Ccl6 | 0 | 0 | 0 | 0 | 10.7717 | 10.2 | 11.2214 | 11.2521 |
| Q8BLB7 | L3mbtl3 | 0 | 0 | 0 | 0 | 10.7065 | 10.6745 | 10.7304 | 10.858 |
| Q9JKE2 | Trem1 | 0 | 0 | 0 | 0 | 10.5613 | 10.7509 | 10.9611 | 10.9824 |
| P33434 | Mmp2 | 0 | 0 | 0 | 0 | 10.3813 | 11.1978 | 10.8481 | 11.1757 |
| Q3U0L2 | Ankrd33b | 0 | 0 | 0 | 0 | 10.2615 | 10.051 | 10.5372 | 10.4443 |
| Q6AW69 | Cgnl1 | 0 | 0 | 0 | 0 | 10.1325 | 10.3246 | 10.0921 | 10.1673 |
